# ILnc: Prioritizing Long Non-coding RNAs for Pan-cancer Analysis of Immune Cell Infiltration

**DOI:** 10.1101/2022.03.10.483725

**Authors:** Xinhui Li, Changbo Yang, Jing Bai, Yunjin Xie, Mengjia Xu, Hui Liu, Tingting Shao, Juan Xu, Xia Li

**Affiliations:** College of Bioinformatics Science and Technology, Harbin Medical University, Harbin 150081, China; Key Laboratory of Tropical Translational Medicine of Ministry of Education, College of Biomedical Information and Engineering, Hainan Medical University, Haikou 571199, China

**Author notes:** Equal contribution (Xinhui Li, Changbo Yang, Jing Bai). Corresponding authors. (Li X), (Xu J), (Shao T).

**Keywords:** Long non-coding RNA, Pan-cancer, Immune infiltration, Prognosis

## Abstract

The distribution and extent of immune cell infiltration into solid tumors play pivotal roles in cancer immunology and therapy. Here we introduced an immune long non-coding RNA (lncRNA) signature-based method (ILnc), for estimating the abundance of 14 immune cell types from lncRNA transcriptome data. Performance evaluation through pure immune cell data shows that our lncRNA signature sets can be more accurate than protein-coding gene signatures. We found that lncRNA signatures are significantly enriched to immune functions and pathways, such as immune response and T cell activation. In addition, the expression of these lncRNAs is significantly correlated with expression of marker genes in corresponding immune cells. Application of ILnc in 33 cancer types provides a global view of immune infiltration across cancers and we found that the abundance of most immune cells is significantly associated with patient clinical signatures. Finally, we identified six immune subtypes spanning cancer tissue types which were characterized by differences in immune cell infiltration, homologous recombination deficiency (HRD), expression of immune checkpoint genes, and prognosis. Altogether, these results demonstrate that ILnc is a powerful and exhibits broad utility for cancer researchers to estimate tumor immune infiltration, which will be a valuable tool for precise classification and clinical prediction.

## Introduction

Immune cells, including innate immune cells [*e*.*g*., macrophages, neutrophils, dendritic cells (DC), and nature killer (NK) cells] and adaptive immune cells (*e*.*g*., T and B cells), are important components of the immune system. Malfunctions of immune cells are always associated with diseases, including cancers [1,2]. Thus, investigating immune cell distribution in cancers could provide important insights into immune status, cancer progression, prognosis, as well as immunotherapy [3]. In addition, immunotherapy has obtained great successes in the clinical application, particularly in the field of immune checkpoint blockade [4]. The same tumor type has different immunotherapy effects due to the different tumor-infiltrating lymphocytes in the tumor microenvironment [4]. For instance, metastatic melanoma patients with different immune cell compositions respond differently to anti-PD-1 treatment [5]. The level of tumor-infiltrating lymphocytes in the tumor microenvironment directly affects the progress of tumor and treatment effect [6-8]. Therefore, identifying the composition of immune cells and exploring the dynamic changes in the proportion of immune cells during tumor progression will help to provide personalized immunotherapy for patients.

Long non-coding RNAs (lncRNAs), more than 200 nt in length, are one of the major classes of regulatory RNAs that have been widely investigated, playing crucial roles in diverse cellular processes from normal development to disease progression [1,9,10]. The extensive changes in lncRNAs expression during immune cell development, differentiation and activation affect the development of cancer. For example, *Inc-Tim3* is a marker of CD8^+^ T cells and it can induce nuclear translocation of *Bat3* by binding with *Tim3* to promote CD8^+^ T cell exhaustion in hepatocellular carcinoma [11]. *LINC00092* is specifically expressed in cancer-associated fibroblasts (CAFs) from ovarian serous cystadenocarcinoma (OV) microenvironment and causes poor prognosis [12]. *Lnc-BM* and *SNHG1* respectively have been found to be markers of tumor-associated macrophages (TAMs) and regulatory T cells (Tregs) in breast cancer microenvironment. *Lnc-BM* can induce the expression of *CCL2* to recruit macrophages [13], while lncRNA *SNHG1* can promote the differentiation of Treg and further promote the malignancy of tumors [14]. The expression level of specific lncRNAs in immune cells could reflect the proportion of corresponding immune cells to a certain extent. Therefore, it is very valuable to explore the lncRNAs characterizing various immune cells and further predict the infiltration of immune cells based on these lncRNAs.

In recent years, several algorithms have been developed to predict the infiltration level of immune cells in the tumor microenvironment, most of which are designed based on protein coding genes. The transcription levels of two genes, *GZMA* and *PRF1*, were used for the first time in an innovative study to assess the cytolytic activity (CYT) of effector cells to elucidate the possible mechanisms of immune escape [15]. Immune cell signature genes can also be used to characterize the degree of lymphocyte tumor-infiltration. For example, Newman *et al*. have developed CIBERSORT to estimate the proportion of 22 immune cells by predefining an immune signature matrix [16]. Butte *et al*. have developed the xCell algorithm that can predict the proportion of immune cells in tumor microenvironment based on the signature genes generated from expression profiles collected by different channels [17]. Moreover, Liu *et al*. have generated the algorithm TIMER to evaluate the proportion of six types of immune cells in tumor tissues based on the gene expression profile from The Cancer Genome Atlas (TCGA) [18]. There are also many other algorithms for predicting the level of tumor infiltrating lymphocyte (TIL) in tumor tissue, such as MCP-Counter [19] and EPIC [20]. DNA methylation was also introduced to predict immune cell infiltration [21]. In addition, other deconvolution methods were developed to infer proportions of cell types from bulk transcriptomic data. For instance, the method DeconRNASeq estimates the proportions of distinctive tissue types based on mRNA-Seq data [22]. Recently, several methods have been developed by integrating scRNA-seq data as reference to characterize cell type compositions, such as SCDC [23], MuSiC [24] and DWLS [25]. Francisco Avila Cobos *et al*. comprehensively evaluated deconvolution methods from different aspects and provided general guidelines [26]. However, there is still a lack of methods designed for estimating the abundance of numerous immune cell types based on non-coding RNAs.

In this study, we present an immune lncRNA signature-based method (ILnc), to robustly and precisely estimate the abundance of 14 immune cell types [B cell, CD4^+^T cell (CD4^+^ T), CD8^+^ T cell (CD8^+^ T), Treg, CD3^+^ T cell (CD3^+^ T), DC, monocyte, M0-macrophage (M0), M1-macrophage (M1), M2-macrophage (M2), macrophage, neutrophil, NK cell, and granulocyte] from lncRNA transcriptomic data. We compared ILnc with current prevalent methods for predicting the proportion of immune cells infiltration. Furthermore, immune properties of characteristic lncRNAs of immune cells were systematically characterized. We applied the method to TCGA pan-cancer data to describe the tumor microenvironment and explore the influence of immune cells on clinical progression of patients. Finally, we identified six immune subtypes that encompass multiple cancer types and are hypothesized to define immune response patterns impacting prognosis.

## Methods

### Overview of ILnc

The algorithm of ILnc mainly includes three steps: generating initial lncRNA signatures, scoring lncRNA signatures, and selecting the best lncRNA signatures (**Figure 1**).

**Figure 1.**
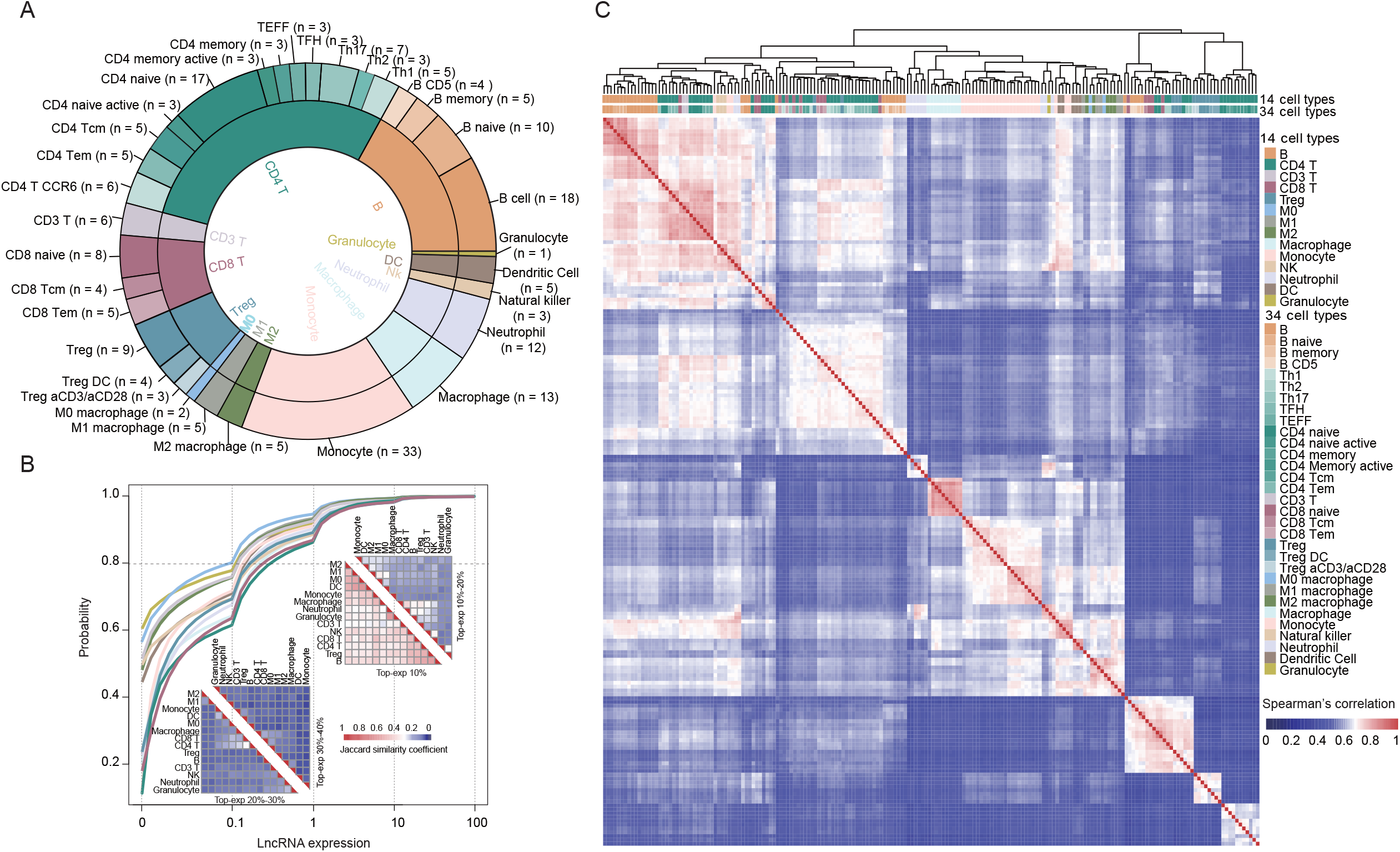
The schematic diagram of ILnc. The three parts are the three main steps of ILnc. The lncRNA, long non-coding RNA expression profiles of 14 immune cells were obtained from GEO, Gene Expression Omnibus database. For each cell, the initial signature lncRNA set was constructed by expression value and scoring every lncRNA by its contribution to estimation accuracy. The best signature lncRNA sets correspond to the largest estimation accuracy and with high variance cross 14 cell types.

#### Generating initial lncRNA signatures

The fragments per kilobase of per million (FPKM) of lncRNA transcriptome of pure immune cells was converted to log scale by adding 0.001 to restrict the inclusion of small changes and followed by log 2 conversion. For each immune cell type *i*, the initial signature sets L _*i*_ (*i* = B, CD4^+^ T, CD8^+^ T, Treg, CD3^+^ T, DC, monocyte, M0, M1, M2, macrophage, neutrophils, NK, granulocytes) were generated by selecting highly expressed lncRNAs, the sum of which transcription levels accounted for 85% of the total transcription levels of all lncRNAs. The initial estimation accuracy *P*_*i*_ of each initial signature set *L*_*i*_ was calculated by the deconvolution method and realized by CIBERSORT based on these selected lncRNA expression profile of pure immune cells.

#### Scoring lncRNA signatures

The contribution of each lncRNA in each initial signature set was evaluated through one by one removing from each initial signature set. The contribution score *S*_*i*_*j*_ of each lncRNA_*i*_*j*_ in the initial signature set *L*_*i*_ was defined as the differences between *P*_*i*_ and *P*_*i*_*j*_.

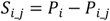

The P_*i* _*j*_ was obtained as the same as *P*_*i*_ after deleting this lncRNA_*i*_*j*_ from the initial signature set *L*_*i*. *n*_*i*_ means the number of lncRNAs in *L*_*i*_ .The higher the *Si*_*j*_ value, the greater the contribution for the lncRNA_*j*_ to the cell type *i*.

#### Selecting the best signatures

For each initial signature set *L*_*i*_, we first sorted all lncRNAs in descending order according to score *S*. Then, the top m lncRNA as the lncRNA set *L*_*i* _*m*_ was used to calculate estimation accuracy *P*_*i* _*m*_ (*m* = 30, 35, 40, 45… *n*_*i*_). LncRNAs with high variance across 14 cell types were identified by analysis of variance as *L*_*v*_ (*FDR* < 0.3). Finally, the lncRNA signature set *L*_*i* _*m*_ with the largest *P*_*i* _*m*_ filtered with *L*_*v*_ was defined as the signature set of the immune cell *i*.

To conveniently use this method, we developed the algorithm into an R package ILnc. ILnc is an independent R package and the output formats of results are similar as other methods, which are to simplify future integrative analysis if necessary.

### Data resource

The 29 RNA-seq datasets of pure immune cells including 184 samples were downloaded from the Gene Expression Omnibus (GEO) database through manual retrieval using keywords “immune cells” and “RNA-seq”. Another 63 RNA-seq datasets were downloaded from Ranzani *V et al*. [23]. The data preprocessing and normalization is similar to previous studies [24,25]. First, raw RNA-seq data were aligned against the human genome hg19 using TopHat2 [26] and assembled into transcripts using cufflinks [27]. The lncRNA expression level was represented by FPKM value which removes the effects of both library size and gene length, and further performed log transformation. Similarly, the transcriptome of protein-coding genes was also constructed. The transcriptomic data of all lncRNAs were screened out based on annotation from the GENCODE consortium. The samples with all zero expression were deleted and the lncRNAs with expression value (> 0) in less than 100 samples were also deleted. All 247 samples are pure immune cell line data of 34 cell types, and immune cell subsets can be further grouped into 14 major leukocyte types on the basis of shared lineage. By calculating the average value of samples of the same type, transcriptomic data of 15,444 lncRNAs corresponding to 14 immune cell types are obtained.

RNA-seq data of 15 immune cell types were obtained from DICE database as a validation dataset and merged into 7 types. The RNA-seq data of TCGA samples and the clinical information were downloaded from Broad GDAC Firehose. Moreover, the lncRNA expression profiles of 272 glioma samples were obtained from GEO (GSE48865), including 100 grade II, 72 grade III, and 100 grade IV samples [28,29]. For each immune cell type, idealized mixture was constructed by combining a certain percentage of the purified immune cell with other types of immune cells by random selection and the mean of the expression of these cells was used as the expression of one mixed sample.

### Immune related functional analysis of lncRNA signature

We analyzed functions of 14 lncRNA signature sets using Kyoto Encyclopedia of Genes and Genomes (KEGG) and Gene Ontology (GO) terms enrichment analysis based on corresponding co-expressed protein-coding gene sets. For each lncRNA signature set, if the expression of one protein-coding mRNA is significantly correlated with more than 10% lncRNAs in this set, this protein-coding gene is considered to be co-expressed (|r| ≥ 0.6). The hypergeometric test was used to identify the significantly overrepresented immune pathways of a specific co-expressed protein-coding gene sets based on the ImmPort database [30]. Known marker genes of immune cells were obtained from the CellMarker database including B cells, granulocytes, T cells, macrophages, monocytes, dendritic cells, neutrophils, NK cells and regulatory T cells [31]. For each type of immune cells, we sorted all lncRNAs based on the median Pearson correlation coefficient between a lncRNA and all marker genes, and the samples used in the calculation are limited to this type of immune cells. Gene set enrichment analysis (GSEA) was used to estimate whether co-expressed genes with signature lncRNAs were overrepresented in known marker genes of immune cells.

### Identification of six immune subtypes

We identified six immune subtypes based on the infiltration level of immune cell of cancer patients using hierarchical clustering in the pheatmap package of R program. The package used popular clustering distance (“euclidean”) and method (“complete”) implemented in dist and hclust functions in R. Here, two immune cell types with very low infiltration level in most samples were eliminated.

We also analyzed the differences in homologous recombination deficiency (HRD) score [32], CYT score [33], major histocompatibility complex (MHC) [34], immune score [35] and survival time among six subtypes.

### Statistical Analysis

Basic statistical analyses, such as the Wilcoxon rank-sum test and Pearson correlation coefficient, were performed using R program. Cox regression, log-rank test, and Kaplan-Meier in R package “survival” were used to assess the relationships between the proportion of immune cells and survival time. The p value threshold was 0.01 in case studies.

## Results

### Landscape of lncRNA transcriptome in human immune cells

To assess lncRNA expression in human immune cells, the original 30 RNA-seq datasets of pure immune cells including 247 samples were obtained from the GEO and PubMed database through manual retrieval. These immune cells already have been with a defined cell type (Table S1). The lncRNA transcriptional profiles of 34 immune cell subsets were constructed based on these datasets using TopHat and Cufflinks, which almost cover most types of immune cells (**Figure 2**A). We further filtered samples and lncRNAs with low expression (see Methods). In addition, we performed bench correction by using the ComBat function in R language, and found a strong positive correlation between data before and after batch correction (R = 0.99). Finally, we obtained a transcriptional map of 15,444 lncRNAs from 218 samples. In order to analyze lncRNAs in immune cells more systematically, immune cell subsets can be further grouped into 14 major leukocyte types on the basis of shared lineage (B cell, CD4^+^ T, CD8^+^T, Treg, CD3^+^ T, DC, monocyte, M0, M1, M2, macrophage, neutrophil, NK, and granulocyte) to refer to immune cell function (Figure 2A).

**Figure 2.**
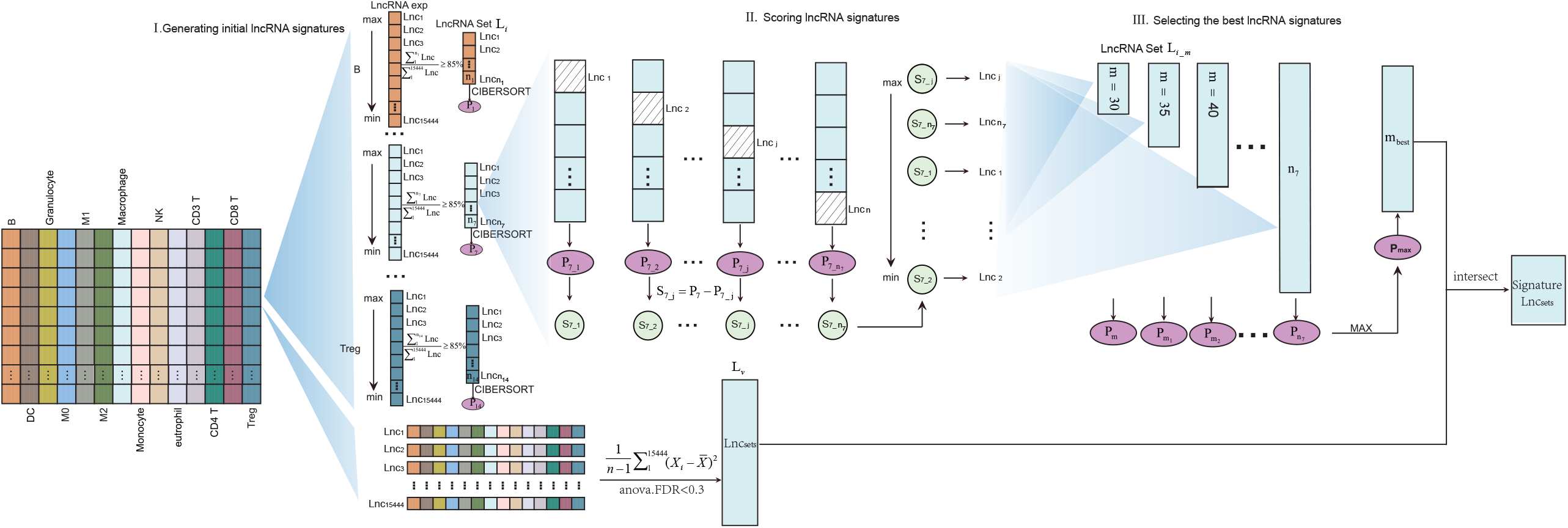
Transcriptome of lncRNAs in immune cells. A. A summary of the data used in the study to generate the lncRNA signatures. The outer circle shows the 34 pure cell types and the number of samples of the cell types, the inner circle shows the number of samples for 14 immune cell types. **B**. Hierarchical cluster analysis of correlations between samples based on lncRNA transcription data. Different legends represent different sample classifications. **C**. The cumulative probability in different expression intervals and Jaccard similarity coefficients between lncRNA sets by 10% threshold.

Next, we evaluated the expression level of these lncRNAs in 14 immune cell types. We found the proportion of lncRNAs, expression of which are less than 0.1, is more than 60%. The proportion of lncRNAs with expression values greater than 10 accounted for less than 2% (Figure 2B). We ranked all lncRNAs for each immune cell type by expression in descending order and equally divided them into 10 groups. The Jaccard similarity coefficients were calculated between each cell type for each group and found the shared lncRNAs among immune cells was a few in each group (Figure 2B and Figure S1). These results suggested that most lncRNAs have lower expression values in immune cell samples, and although most lncRNAs were shared among immune cells, the expression level of lncRNAs was different. As lncRNAs are generally more tissue-specific than protein-coding genes [36], we assessed the immune cell subgroups specificity of lncRNAs by Hierarchical clustering (Figure 2C and Figure S2). Then, we found that lncRNAs showed similar expression patterns within subgroups, and immune cell subtypes with the same origin are clustered together. For example, CD3^+^ T cells, CD4^+^ T cells, and CD8^+^ T cells which all belong to T cells tend to be clustered together and generation of them all occur in the thymus from precursors arriving from the bone marrow [37]. M0, M1, and M2 cells which all belong to macrophages are clustered to a category, and they all develop from monocytes [38]. Altogether, based on RNA-seq analyses of highly purified immune cell subsets, we provided a comprehensive landscape of lncRNA expression in human immune cells. The results suggested that lncRNA expression is specific in immune cells and may have the potential to predict fractions of multiple immune cell types in lncRNA expression profiles of admixtures.

### Prioritizing lncRNA signatures of immune cells is the key of ILnc

We developed a lncRNA-based deconvolution pipeline for application in immune cell infiltration, and the creation of lncRNA signatures is critical. We interrogated the lncRNA transcriptome dataset for the presence of lncRNA signatures in the 14 immune cell subgroups. We proposed a multi-step method to identify the lncRNA signature sets for 14 immune cell subgroups based on the lncRNA transcription profile, which is the key to predict the immune cell infiltration. The hypothesis of our method ILnc is that lncRNAs with the greatest contribution to each cell type might be the important component of lncRNA signatures, because there are overlap among known signature sets of different cell types. The multi-step method includes three parts (Figure 1 and Methods). First, the highly expressed lncRNAs were served as the initial signature sets the sum of expression values of which accounted for 85% of the total expression values in each cell subtype in descending order. Next, the contribution score of each lncRNA was evaluated based on one by one removed from each initial signature set. The lncRNAs were sorted according to the contribution rate in a descending order, and the estimation accuracy of each immune cell subgroup was calculated based on the top lncRNAs as candidate signature sets with different thresholds. The estimation accuracy of each immune cell subgroup represents the percentage of immune cells predicted by pure immune cells. We found that the estimation accuracy became higher with the increase of the number of lncRNAs and soon reached a stable level (**Figure 3A**). We calculated the variance of these lncRNAs across all pure cells, and found that signature lncRNA candidates tended to be highly variable (Figure S3). Thus, the final lncRNA signature set was further filtered with high variance. Lastly, 14 lncRNA signature sets for each immune cell subgroup were constructed, which distinguish human immune cell phenotypes. We also found that the size of lncRNA characteristic sets varied in a wide range from 189 to 895 (Table S2).

**Figure 3.**
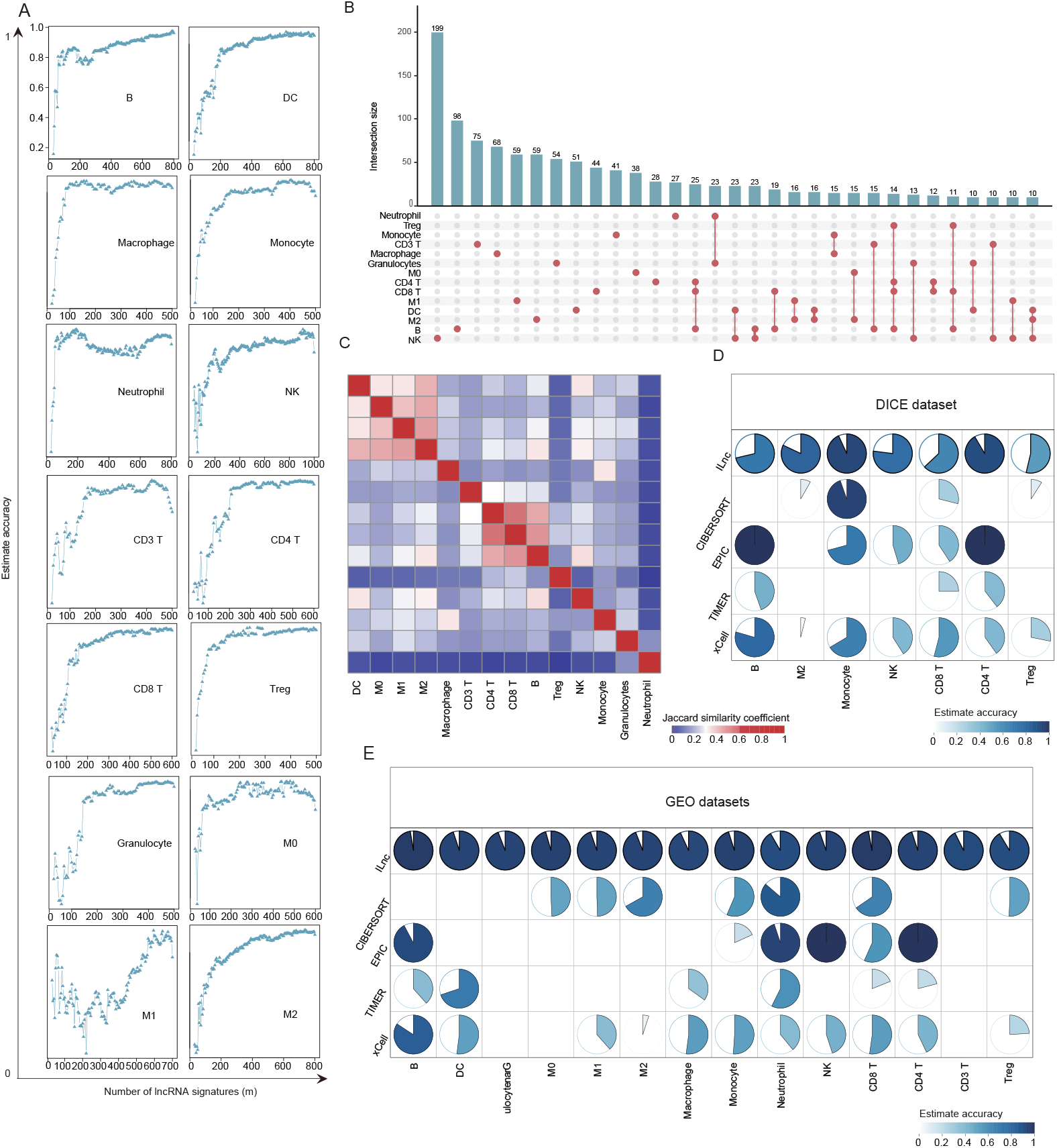
Constructing and characterizing the lncRNA signature sets. A. Estimated results of multiple high-score lncRNA sets during the process of “choosing the best signature”. **B**. The overlap between signature sets was shown by the upset graph, and the threshold is 10. **C**. The bidirectional clustering graph shows the jaccard similarity coefficients between the 14 signature sets. D. Accuracy comparison of the ILnc and another four methods. E. Accuracy comparison of the ILnc and another four methods based on external data sets. The lncRNA expression profile and lncRNA signatures were used in ILnc and the mRNA expression profile and mRNA signatures were used in running another four methods (CIBERSORT, EPIC, TIMER, and xCell) in their default parameters.

To further explore the association among the 14 signature sets of immune cells, we firstly described the distribution of characteristic lncRNAs across immune cell signature sets by calculating the number of signature sets in which each lncRNA is involved. We found that lncRNAs have different relative contributions to distinguish human immune cell phenotypes, and the large majority of lncRNAs occur in more than one immune cell subgroups (Figure S4). Furthermore, approximately 3.4% to 22.2% of the lncRNAs were involved in one signature set across immune cell characteristic lncRNA sets and most of them were presented in NK cell (199/22.2%), CD3^+^ T cell (75/18.7%) and macrophage (68/16%), representing approximately 38.82% of all the characteristic lncRNAs (Figure 3B). Conversely, the fraction of lncRNAs shared by at least 5 signature sets ranged from 3.8% to 19.8%, and represented only approximately 25.33% of all the characteristic lncRNAs. For example, lncRNA *NIFK-AS1* was involved in 5 immune cell sets, which suppresses the M2 macrophages polarization by inhibiting *miR-146a* [39]. LncRNA *SNHG1* was involved in 11 immune cell signature sets, knockdown of which will suppress Treg differentiation by increasing the expression of *miR-448* and reducing level of *IDO* [14]. These results suggested that although 75 core lncRNAs occur in most signature sets (> 10), different expression patterns and lncRNA combinations may be used to explain the efficiency to distinguish human immune cell phenotypes. Then we counted the number of lncRNAs specifically shared among different lncRNA sets and found that lncRNAs tend to exist in only one set, and the number of lncRNAs that specifically exist in multiple sets is small (Figure 3B). Further, we calculated Jaccard coefficients between any two signature sets and found that CD4^+^ T cells shares more characteristic lncRNAs with CD8^+^ T cells (Figure 3C). There are 366 common lncRNAs for the two immune cells and 28 lncRNAs are specific. The expression of the common lncRNA is different in two kinds of immune cells (Wilcoxon, *P* = 1.86*e*^−4^). There are 245 common lncRNAs among M0, M1, M2 macrophage immune cells and 156 lncRNAs are specific, and these common lncRNAs also show different patterns of expression in three immune cell types (Figure S5). The results suggested that immune cells with similar origin have more similar lncRNA characteristic sets.

To validate prediction effect of the lncRNA signatures, we applied it to predict external data sets of variably purified immune cell subsets. This test dataset was obtained from the Database of Immune Cell Expression (DICE) database [40], and merged into seven kinds of immune cells subgroups according to the same principle. We found that the results matched ground-truth phenotypes in average 76.0% of these seven immune cell types data sets, and the highest estimation accuracy reached 93.1% for Monocyte (Figure 3D). In addition, we compared the results predicted by the ILnc with four common used methods, including CIBERSORT [16], EPIC [20], TIMER [18], and xCell [17] (Figure 3D and 3E). Because these four methods were developed based on mRNA transcriptome, we used the mRNA expression profile constructed from the same dataset as lncRNAs and their separate mRNA signatures with default parameters. We found that the abundance of immune cells estimated by ILnc is robust. To further evaluate their performance, we constructed a simulated mixture population dataset with well-defined composition for each type of immune cell. ILnc was also used on the mixed population data. As a result, significantly high concordances were discovered between the predicted percentage and the true percentage in most immune cells (Figure S6). All results suggested that ILnc is robust and able to accurately estimate the abundance of 14 immune cell types.

### Associations of lncRNA signatures in ILnc and immune-related functions

More and more studies have shown that lncRNAs are involved in immune biological processes, including regulating the homeostasis of immune cells and inflammatory response [41,42]. Therefore, we reasoned that if these characteristic lncRNAs contribute to the classification of immune cells, and they would be more likely to participate in functions of corresponding immune cells and co-express with known marker genes of immune cells. Firstly, the biological functions of characteristic lncRNAs were inferred on the basis of co-expressed protein coding genes (see Methods). The co-expressed gene set of each signature set was obtained in which the Pearson correlation coefficients of protein coding genes are greater than 0.6 with at least 10% lncRNAs. Functional enrichment analysis was performed on co-expressed coding genes (Table S3). We found that these characteristic lncRNAs are significantly enriched to immune-related functions. For example, most of the lncRNA signature sets were significantly participated in immune response functions and inflammatory response (**Figure 4A**). Particularly, we found several characteristic lncRNAs are involved in the regulation of functions consistent with the immune cell type they represent. KEGG pathways enrichment analysis was also performed on co-expressed coding genes (Table S4). The CD4^+^ T signature lncRNA set is enriched in the T helper cell differentiation pathway, and the signature lncRNA set of CD8^+^ T cells and Tregs are enriched in *CTLA4* signaling in cytotoxic T lymphocytes pathway (Figure 4B). Studies have shown that lncRNAs are involved in the key biological processes of CD4^+^ T cell response in infection [43] and regulate immune response such as *NKILA* regulates T cell sensitivity to Activation-induced cell death (AICD) by inhibiting *NF-κB* activity [44]. Additionally, we found that the characteristic lncRNAs of 14 immune cell types significantly participate in 17 immune-related pathways from the ImmPort database [30] (Figure S7). For example, the B cell signature lncRNAs are enriched in B cell receptor signaling. These results suggest that the characteristic lncRNAs identified by us contribute to immune-related functions.

**Figure 4.**
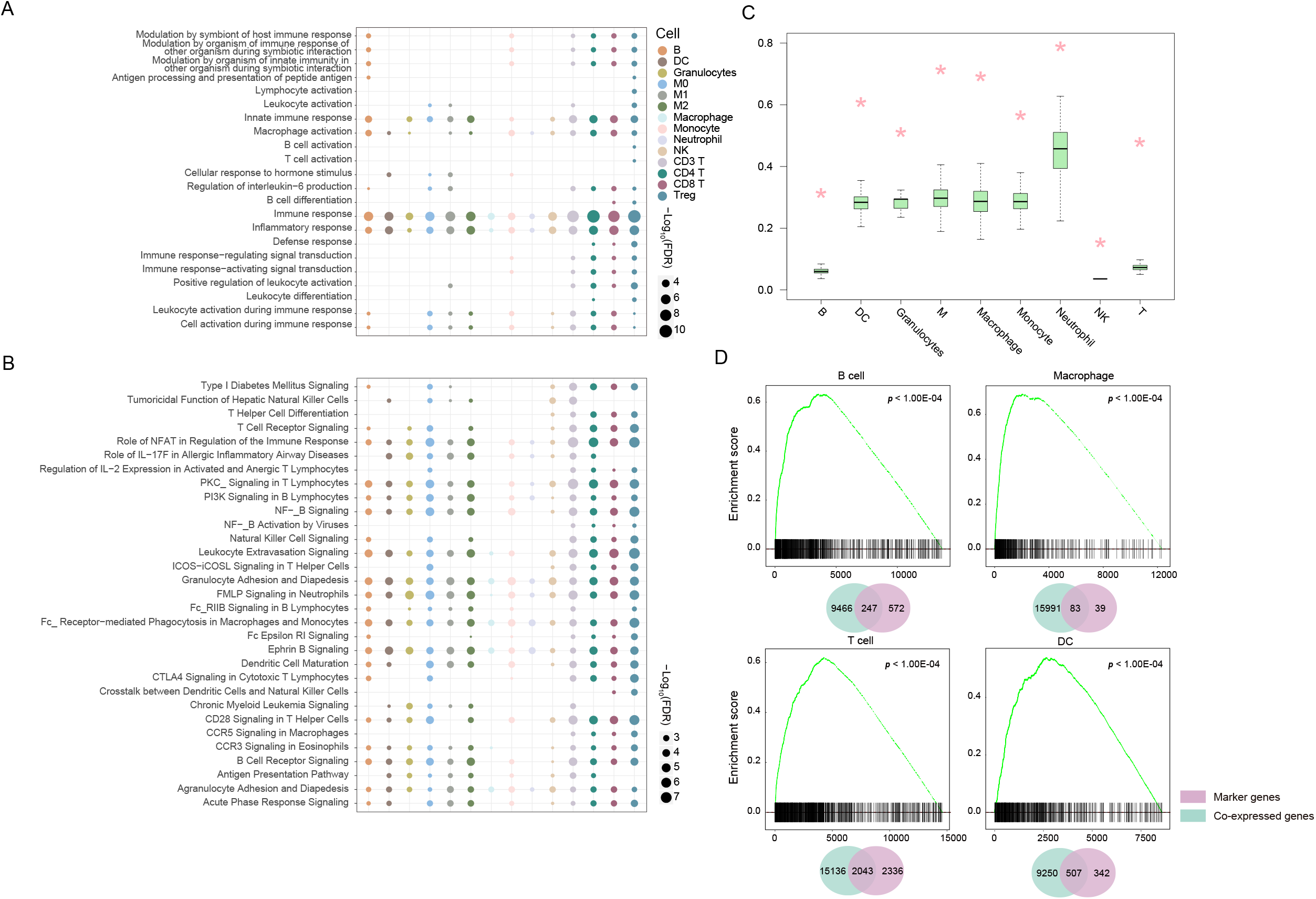
LncRNA signature sets related to immune functions. A. Bubble chart shows the functional enrichment of the co-expressed gene sets of 14 immune cell lncRNA signature sets in GO, Gene Ontology database. **B**. Bubble chart shows the functional enrichment of the co-expressed gene sets of 14 immune cell lncRNA signature sets in KEGG , Kyoto Encyclopedia of Genes and Genomes pathways. **C**. Pink asterisks indicate the proportion of co-expressed gene sets (9 lncRNA signature sets) in the marker gene set. The boxes represent the proportion corresponding to 500 random non-signature sets. **D**. The lncRNAs of the immune cell signature sets is significantly enriched in the immune cell marker genes drawn by the GSEA , Gene set enrichment analysis package. The Venn diagram shows the overlapping numbers of co-expressed gene sets and marker gene sets.

We further verified the characteristic lncRNAs based on well-studied protein coding gene markers of immune cells. We found that co-expressed genes of the lncRNA signature sets have more overlap with marker genes of corresponding immune cell than random lncRNA sets (Figure 4C). For example, 77.66% of known protein coding gene markers of neutrophils are co-expressed with the characteristic lncRNAs of neutrophils, significantly higher than non-characteristic lncRNAs. For each lncRNA signature set, we next sorted all genes according to the mean of correlation coefficients with characteristic lncRNAs, and found that marker genes are significantly enriched in the high correlation group by using GSEA (*P* < 1.00*e*^−4^, Figure 4D). The lncRNA *Flicr* has been found to modulate the expression of marker gene *FOXP3* and enhance the immunosuppressive function of Tregs [45]. We found that the lncRNA signature sets of B cells, DC, macrophages, and T cells are significantly high correlated with the marker genes (Figure 4D). These results suggested that the lncRNA signature sets are closely related to immune functions and marker genes of 14 types of immune cells, and the predictive potential of characteristic lncRNAs is further confirmed from functional level.

### Clinical relevance of immune cell infiltration predicted by ILnc across cancer types

Increasing evidences have shown that the occurrence and development of cancer are closely related to the composition of immune cells in cancer and the infiltration of different immune cells could dramatically influence the treatment strategy and prognosis [8]. To demonstrate the application of lncRNA signature sets for cancer research, we analyzed 10363 samples of 33 cancer types from TCGA with lncRNA expression data to survey the difference of immune cells infiltration (Figure S8-S10). We found that the infiltration levels of macrophages and CD4^+^ T cells are high in most cancer types (**Figure 5A**). Macrophages are crucial drivers of promoting inflammation in tumor and TAMs make the tumor microenvironment conducive to tumor progression and metastasis [46]. Additionally, we found different cancer types have different levels of immune cell infiltration. For example, the infiltration level of B cell is the highest in lymphoid neoplasm diffuse large B-cell lymphoma (DLBC) and the infiltration level of most immune cells is low in lower grade glioma (LGG) [47,48]. Several studies have found that the infiltration levels of immune cell are always relatively low in glioma [49,50]. Tumor tissues have a diverse mixture of tumor and non-tumor cells within their microenvironments [51]. The tumor purity quantifying tumor cell content in the tissue generally is negatively correlated with the proportion of immune cells infiltration [52]. To indirectly confirm immune cells infiltration predicted by us, we calculated the correlation between the infiltration proportion of various immune cells with tumor purity in the 33 tumor types. At the pan-cancer level, about infiltration level of 80% of immune cells were significantly negatively correlated with tumor purity in 33 cancers (Figure 5B). The infiltration proportion of macrophage is significantly negatively correlated with tumor purity in most tumors, including lung squamous cell carcinoma (LUSC) and thyroid carcinoma (THCA). Additionally, we also respectively calculated correlation coefficients between the level of immune cell infiltration predicted by us and the results obtained from the xCell method for 14 immune cells in each cancer [17]. We found that most of our predicted results are significantly positively correlated with the results of xCell (Figure S11). These results further strengthened that the lncRNA signature sets could be used to predict immune cell infiltration.

**Figure 5.**
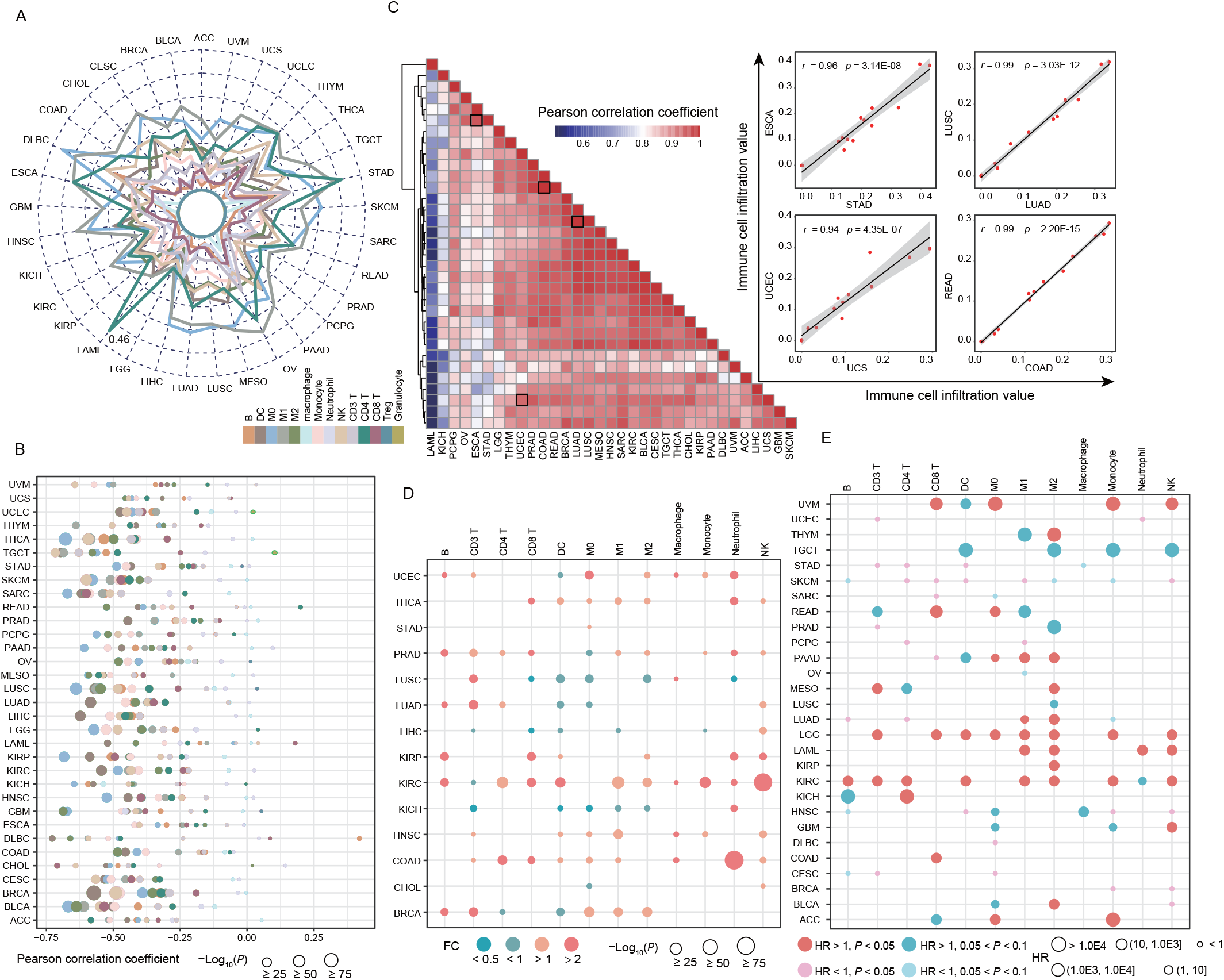
The application of lncRNA signature set in pan-cancer. A. The radar chart shows the infiltration of 14 immune cells in 33 cancer types. **B**. The bubble chart shows the correlation between the infiltration of immune cells and the tumor purity in the pan-cancer. **C**. The bidirectional clustering diagram shows the similarity of immune infiltration between 33 cancers. The scatter plots on the right shows the similarities between several cancers. **D**. The size of the dots indicates the significance of difference of immune cell infiltration between cancer samples and normal samples of 14 cancer types. Red bar indicates more immune cells infiltrating in cancer samples, and blue bar indicates more infiltration in normal samples. **E**. The correlation between immune cell infiltration and prognosis of cancer patients was analyzed by cox-risk regression. The size of the dots represents the HR, Hazard ratio value, and the color represents the p value.

Next, we hypothesized that if cancer types exhibit more similar the immune cell infiltration patterns, they are more likely to be with similar immune microenvironment. We computed a paired similarity based on the infiltration level of 14 immune cells and found that some cancers showed greater similarity to each other, such as lung adenocarcinoma (LUAD), LUSC, bladder urothelial carcinoma (BLCA), cervical squamous cell carcinoma and endocervical adenocarcinoma (CESC), head and neck squamous cell carcinoma (HNSC), breast invasive carcinoma (BRCA), and kidney renal clear cell carcinoma (KIRC), in which a general immune related trend has emerged (Figure 5C). Especially some cancer types with similar tissue origin, such as LUAD and LUSC, rectum adenocarcinoma (READ) and colon adenocarcinoma (COAD), were closely clustered together and the correlation coefficient of immune cell infiltration was as high as 0.99 [53,54].

To study the distribution of infiltrating immune cells in the tumor and adjacent tissues, we focused on 14 cancer types for which the lncRNA expression profiles of adjacent tissues were available. We predicted the level of immune cell infiltration in tumors and adjacent tissues respectively using ILnc (Figure S10), and found that the infiltration level of many immune cell types was significantly different between tumors and adjacent tissues in most cancers (Figure 5D). The M2 macrophage was markedly enriched in the nidus for half of all cancer types, which is consistent with their characteristics of immunosuppression [55]. In contrast, DC showed higher infiltration in adjacent tissues of four cancers [LUSC, LUAD, liver hepatocellular carcinoma (LIHC), kidney chromophobe (KICH), and BRCA]. In addition, we also found that the proportion of some immune cells infiltrating in tumor tissues was significantly lower than in adjacent tissues of LUSC, which is supported by previous observations showing that fractions of some immune cells were significantly lower in tumor tissues than normal tissues in lung cancer, such as DC, monocyte, M0 macrophage and CD8^+^ T cell [56]. In addition, we also found that the proportion of most immune cells infiltrating in tumor tissues was significantly higher than in adjacent tissues in KIRC.

Immune cells infiltration have been linked to patient survival in some cancer types, such as DC and NK cell associated with favorable outcomes in skin cutaneous melanoma (SKCM) [57]. We next investigated the effects of immune cell infiltration on clinical outcomes in pan-cancer using cox regression analysis. We identified many significant associations between infiltration of immune cell types and the overall survival of patients in different cancer types (Figure 5E). We found that infiltration of M2 macrophage consistently predicts poor outcome as a risk factor in nearly half of all cancer types, and other immune cell types have cancer-specific effects on prognosis. Studies have found that M2 macrophage can promote colon cancer cell invasion and pancreatic cancer cell metastasis [58,59]. Five kinds of immune cell types were significantly associated with patient survival in SKCM including DCs, NK cells, CD4^+^ T cells, and CD8^+^ T cells, all of which have positive effects on long-term survival in patients (Figure 5E). These results may partially explain the better outcomes of some patients undergoing immunotherapy [60]. Furthermore, the infiltration of most immune cells had opposite effects on survival in LGG and KIRC [61]. It has been found overall TIL abundance is associated with shorter survival in LGG and KIRC and it can be explained by the danger of inflammation in the brain and by the role of macrophages in suppressive signaling [61]. We also found that the same immune cells have different effects on survival in different cancer types from the same tissue. For example, the infiltration of most immune cells was significantly associated with survival in KIRC, but only M2 was significantly associated with survival in kidney renal papillary cell carcinoma (KIRP), B cells and CD4+ T cells were risk factors in KICH. We further compared the performance of our method with other methods in tumor survival and outcome. We found some consistent results obtained by three different methods (Figure S12). For example, the infiltration of M2 predicts poor outcome, and is a risk factor in most of cancer types. In addition, we found that neutrophil infiltration is a risk factor for KIRC by using our method, but the other two methods did not predict this association. This result indicates that lncRNA might be related with the functional role of Neutrophil infiltration in KIRC. In addition, the 272 glioma samples, covering patients in stages 2, 3, and 4, were estimated immune cell infiltration proportions by ILnc. The results showed that as the malignant degree of glioma increased, the proportion level of CD8^+^ T cells decreased and neutrophile cells increased (Figure S13). The results suggest that the lncRNA-based immune cell infiltration prediction method developed by us could comprehensively predict the infiltration level of immune cell in the tumor and assist to excavate the prognostic immune cell markers. In addition, we further analyzed the association of signature lncRNAs with cancer stage and treatment outcomes. We found that the expression levels of some lncRNA signatures were significantly different in different cancer stages and different treatment outcomes (Figure S14). For example, the two lncRNA signatures of CD8^+^ T cells, *RP11-255H23*, and *RP11-311H10*, are significantly correlated with the therapeutic effect in KIRC, and the two lncRNAs are highly expressed in patients with complete or partial remission. *GABPB1-AS1* as signature of M2, is significantly lowly expressed in advanced LUSC. These results suggest that signature lncRNAs can be used as a potential marker for stage and treatment outcomes.

### Application of ILnc in cancer immune subtyping

To characterize intra-tumoral immune microenvironment states, we identified the cancer immune subtypes based on the infiltration level of 12 immune cells using cluster analysis for 33 cancer types. The six cancer immune subtypes C1-C6 (with 4161, 3210, 535, 207, 230, and 1778 samples respectively) were characterized by a distinct immune cell distribution (**Figure 6**A) and showed some distribution characteristics of cancer and immune indicators in these six subtypes of samples (Figure 6B-C). In C5 subtype, the infiltration level of most of immune cells tends to be high. OV and stomach adenocarcinoma (STAD) are enriched in C5 [62, 63]. C5 contains a relatively small number of cancer samples and this is consistent with the general finding of immunosuppression in cancer tissue [64-66]. C2 is enriched in LGG, prostate adenocarcinoma (PRAD) and pheochromocytoma and paraganglioma (PCPG) and has a lower level of macrophages than other subtypes. C1 has higher levels of M0 macrophages. HNSC, LUSC, and KIRC are enriched in C1. The C3 group has a higher level of B cell and CD4^+^ T cell infiltration and OV are enriched in C3. C4 has higher levels of DC, M0 macrophage, M1 macrophage and CD4^+^ T cell infiltration. Further, to characterize the biological characteristics of each cluster, functional enrichment analysis was performed for each cluster based on these differentially expressed genes. We found that C6 is significantly enriched in regulation of granulocyte macrophage colony-stimulating factor production and regulation of natural killer cell activation. This result is consistent with the distribution of immune cell infiltration in C6, which has higher levels of macrophages and NK cells. C1 is significantly enriched in metabolism-related process, and C4 is significantly enriched in functions related to development and differentiation. Many receptor signaling pathways are significantly enriched in C5, such as receptor signaling pathway via *JAK-STAT*, phospholipase C-activating G protein-coupled receptor signaling pathway and receptor signaling pathway via *STAT*. C3 is significantly enriched in humoral immune response and serotonin receptor signaling pathway function, etc. These results suggested that clusters have different functional characteristics. Immune subtypes identified by us spanned anatomical location and tumor types, while individual tumor types varied substantially in their proportion of immune subtypes. LGG, KICH, PCPG, and PRAD are mainly concentrated in C2 immune subtype, while acute myeloid leukemia (LAML) is concentrated in subtype C6. Conversely, some cancers, esophageal carcinoma (ESCA), and glioblastoma multiform (GBM), are distributed in multiple subtypes (Figure 6B). We also identified cancer clusters based on the infiltration level or score calculated by CIBERTOSRT or xCell. There are large number of sample clusters overlapping with those identified by the two methods, for example, C1 with CC1, C6 with CC6, C1 with XC1, and C6 with XC6 (Figure S15A). We also found a similar distribution of well-known immune markers among these cancer clusters identified by different methods. The patients in C3, CC3, and XC3 exhibit higher HRD scores than other subtypes (Figure S15), and patients in C6 and CC6 clusters had higher immune, CYT, and MHC scores. These distributions of immune cell infiltration represent signatures of the tumor immune microenvironment that largely cut across traditional cancer classifications to create subgroups and suggest future cancer treatment may be independent of histologic type [67].

**Figure 6.**
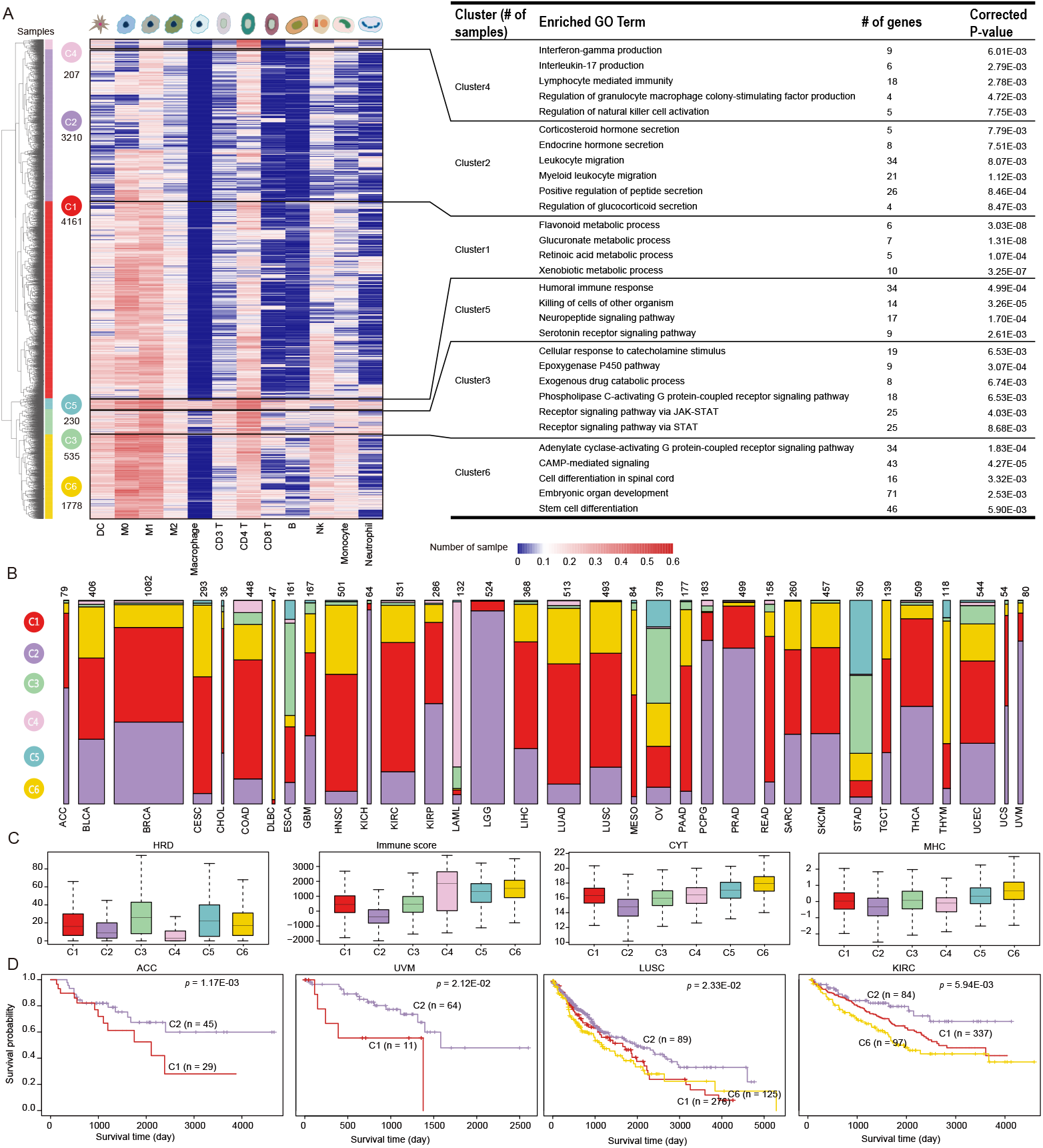
Immune subtypes of pan-cancer. A. The pan-cancer samples were divided into 6 immune subtypes based on immune cell infiltration rate by the hierarchical clustering. For each cluster, the differentially expressed protein coding genes with other 5 clusters were used to functional enrichment analysis. **B**. The stacked map shows the distribution of each cancer sample among the six subtypes. **C**. The differences of five immune related scores among six subtypes. **D**. Survival analysis of four cancers between different clusters.

HRD is emerging as an important biomarker and a potentially effective adjunct to enhance immunogenicity of tumors [32] (Figure 6C). By comparing HRD scores across different subtypes, we found that the distribution of the scores in six subtypes is different. The C3 patients exhibit higher HRD scores than patients in other subtypes. Moreover, immune, CYT and MHC scores remain useful biomarkers for predicting the immune response. We investigated the distribution of the three scores among six immune subtypes and found that C5 and C6 patients had higher immune, CYT and MHC scores than others (Figure 6C). We also explored the expression of immune checkpoint related genes [68] across patients and revealed clear differences between patients in different subtypes (Figure S16). The level of immune cells infiltration in tumor can predict a patient’s clinical outcome in many cancer types [69]. Therefore, we explored the prognostic implications of molecular subgroups. We found that immune subtypes associated with overall survival. The patients in C2 had significantly better overall survival than other patients (Figure S17). Especially in uveal melanoma (UVM), adrenocortical carcinoma (ACC) and LUSC, we found that C1 patients had poorer prognosis than C2 patients (Figure 6D). The overall survival of patients with different subtypes is significantly different in KIRC (Figure 6D). These results suggest that cancer subtypes are remarkable molecular and immunology diversity, which may be helpful to improve personalized cancer immunotherapy.

## Discussion

Changes in immune cell compositions underlie diverse physiological and pathological states. Although there are several methods available for immune cell abundance estimation, most of them are based on protein coding-gene expression. Studies have found that lncRNAs plays considerable regulatory functions in immune cells. In this study, we developed a highly accurate method to estimate the abundance of 14 immune cells based on lncRNA signatures from transcriptomic data, the lncRNA signature sets as core of ILnc, were screened based on the contribution of assessment accuracy, which enhanced the robustness of the prediction. Combining expression of signature lncRNAs with the robust performance of CIBERSORT previously displayed on gene expression allowed us to derive estimation for different infiltrating cell populations. We see ILnc and CIBERSORT as complementary tools for studying the tumor microenvironment in cases where both lncRNA and gene expression data can be obtained. In addition, we applied ILnc to independent data from DICE and found that the estimation accuracy is up to 76.0% on average. Compared with several previous methods, the prediction accuracy of the lncRNA signature sets is better than most prediction methods. At the same time, some signature lncRNAs has been found to be related with immune cells, for example lncRNA *NIFK-AS1* involved in 5 lncRNA signature sets, which can suppress the M2 macrophages polarization by inhibiting *miR-146a* [39]. Collectively, signature lncRNAs could use to estimate the infiltration proportions of immune cells, and provided a new way to study non-coding RNA markers of immune cells. Meanwhile, it still has limitation need to be addressed for our method. In order to reduce the estimation bias caused by the similarity between cell subtypes (CD4^+^ and CD8^+^ T cells, M0, M1, and M2 macrophages), we constructed lncRNA signature sets of 14 immune cell subtypes respectively, and obtained a large number of lncRNA signatures (2187). We will further reduce the number of lncRNA signatures by other scoring algorithms, and optimize the signature sets by more experimentally verified lncRNA markers in the future.

LncRNA expression showed high specificity of tissues and cell types and the extensive changes in lncRNAs expression during immune cell development. Therefore, it is very valuable to explore the lncRNAs characterizing various immune cells and further predict the infiltration of immune cells based on these lncRNAs. At present, some methods based on expression of protein-coding genes or methylation level have been developed to predict the proportion of immune cell infiltration in mixed samples. However, there is a lack of methods based on ncRNAs to predict immune cell infiltration. If only lncRNA expression profiles were detected in mixed samples, lncRNA gene sets could be used to predict immune cell infiltration. When comparing with other methods in our dataset, we found that the performance of our method was better than other methods based on protein-coding genes for some immune cells. Meanwhile, we also combined lncRNA signatures with existing protein-coding gene signatures to compare with lncRNA signatures, and found no significant difference in prediction results between the two signature sets (Figure S18). In future work, we will further optimize the integration of lncRNA and mRNA signatures instead of simple combination to improve predictive effectiveness. Users could choose appropriate signature sets for prediction according to the actual situation of their own dataset and needs.

Consistent with the strong tissue-specificity of lncRNAs, whole lncRNA transcriptome could separate major cell types (Figure 2C). Similarly, the signature lncRNAs were also overlap, indicating their common roles in different cell subtypes. For CD4^+^ T cells and CD8^+^ T cells which contain many cell subtypes, and the overall expression of lncRNAs are different among cell subtypes (Figure 2C). We found that some signature lncRNAs of CD8^+^ T cells were widely expressed in CD8^+^ T cell subtypes. Some signature lncRNAs are specifically expressed in effector memory CD8^+^ T cells, such as *HCG25* and *ARMC2-AS1* (Figure S19). However, the currently publicly available immune cell subtype datas were still limited. We believed that with the increase of pure immune cell data in the future, our method can be extended to a finer scale on immune cell subtypes, especially for CD4^+^ T cells and CD8^+^ T cells.

Increasing evidence suggested that immune cells play critical roles in progression of tumors. A proper proportion of some immune cell subsets, such as Th1, NK, and γδT, could contribute to long-term clinical benefits of anticancer treatments [70,71]. In contrast, the change of the regulatory T cell proportion or their secretions, antigen presentation, and the production of immune suppressive mediators could contribute to cancer immune evasion [72]. Therefore, investigating the abundance of various immune cells could help us understand the tumor immune microenvironment of patient more comprehensively, and benefit to treatment. We applied ILnc to 33 cancers types of TCGA, and found that the heterogeneity of immune infiltration in different cancers and the differences with adjacent tissues are significantly related to the patient’s survival time. And these tumor samples were divided into six immune subtypes by different immune cell infiltration patterns. We found that immune subtypes have significant differences in immune cell infiltration, HRD, expression of immune checkpoint genes and prognosis.

In summary, this study presented an accurate and reliable method to dissect immune cell compositions and explore the infiltration of immune cells in tumor. It may be the first attempt to use lncRNAs to estimate the abundance of multiple immune cells in the transcriptome of cellular heterogeneous tissues such as normal or malignant tissues. ILnc has ability to accurately estimate the abundance of 14 immune cells and reduce estimation bias through individual prediction. The results applied to pan-cancer provide a valuable comprehensive overview for elucidating the interaction between cancer and immunity, and can be used as valuable prognostic predictors to promote the application of cancer immunotherapy and personalized medicine.

## Data availability

R package “ILnc ” developed by us is available in https://ngdc.cncb.ac.cn/biocode/tools/BT007288/releases/1.0 (or https://github.com/hamijia/ILnc). The code used in the work is available in https://ngdc.cncb.ac.cn/biocode/tools/BT007288 (or
https://github.com/hamijia/Code).

## CRediT author statement

**Xinhui Li**: Methodology, Software, Resources, Writing -Original Draft. **Changbo Yang**: Methodology, Resources, Data curation, Validation. **Jing Bai**: Visualization, Methodology, Software. **Yunjin Xie**: Software, Validation, Formal analysis. **Mengjia Xu:** Software. **Hui Liu**: Writing-Reviewing and Editing. **Tingting Shao**: Conceptualization, Supervision, Funding acquisition, Writing-Original Draft, Writing-Reviewing and Editing. **Juan Xu**: Supervision, Conceptualization, Project administration, Funding acquisition. **Xia Li**: Supervision, Conceptualization, Project administration, Funding acquisition.

## Acknowledgments

This work was supported by the National Key R&D Program of China (2018YFC2000100); the National Natural Science Foundation of China (61873075, 32070673, 31871338, 31970646, 62073106, and 32060152); Natural Science Foundation for Distinguished Young Scholars of Heilongjiang Province of China (JQ2019C004) and Natural Science Foundation of Heilongjiang Province of China (LH2020C055).

## Supplementary material

**Figure S1. Jaccard similarity coefficient between high expression lncRNA sets by 10% threshold**.

**Figure S2. The dataset before and after batch correction. A**. Hierarchical clustering graph of Spearman correlation coefficients calculated between samples based on the expression of all RNAs before and after batch correction. **B**. The correlation coefficients between the samples before and after batch correction.

**Figure S3. A. The variance of lncRNA signatures across all pure cells compared with random. B. The significance level of variance analysis of lncRNA signatures across all pure cells compared with random**.

**Figure S4. The stacked graph shows the distribution ratio of lncRNAs in each signature set**. Different colors indicate lncRNAs involved in different numbers of signature sets.

**Figure S5. The expression differentiation of 366 lncRNAs that common for the CD4**^+^ **T and CD8**^+^ **T immune cells, and the 245 lncRNAs common among M0, M1 and M2 macrophage immune cells**.

**Figure S6. The Pearson correlation coefficients between the predicted percentage by our method and the true percentage in the constructed mixed immune population data**.

**Figure S7. The 14 co-expressed gene sets significantly enriched in 13 immune-related pathways from the Immport database by hypergeometry method**.

**Figure S8. The circos diagram shows in detail the immune infiltration profile of 11 cell types in pan-cancer samples**.

**Figure S9. The infiltration of 12 cell types in cancer samples of 33 cancer types and sort cancers by mean value**.

**Figure S10. Comparison of the infiltration of 12 types immune cells between cancer and normal samples in 14 cancer types**.

**Figure S11. The Pearson correlation coefficients between the estimated infiltration of immune cells by our method and xCell**.

**Figure S12.The association between immune cells infiltration predicted by two methods and survival in cancers. A. The association between immune cells infiltration predicted by xCell and survival of samples in 28 cancer types. B. The association between immune cells infiltration predicted by CIBERSORT and survival of samples in 28 cancer types**.

**Figure S13. Independent data from GEO analyzed the relation between stage of glioma patients related to immune cell infiltration**.

**Figure S14. The association between the expression of lncRNA signatures and treatment outcomes or cancer stages of cancer types in TCGA, The Cancer Genome Atlas. A**. The integer shows the number of lncRNA signatures whose expression is significantly related to treatment outcomes in each cancer type based on variance analysis. **B**. The integer shows the number of lncRNA signatures whose expression is significantly related to cancer stages in each cancer type based on variance analysis. The color legend shows the percentage of (A) treatment-related or (B) stage-related lncRNA signatures. **C**. Two examples are given to illustrate the treatment-related lncRNA signatures. **D**. Two examples are given to illustrate the stage-related lncRNA signatures.

**Figure S15. The comparison between ILnc with CIBERSORT or xCell in cancer subtypes. A**. The integer in these figures shows the number of the intersection of samples belonging to different clusters identified by our method with samples of different clusters identified by CIBERSORT. **B**. The distribution of four immune related scores in six immune subtypes identified by CIBERSORT. **C**. The differences in overall survival of samples among six immune subtypes identified by CIBERSORT. **D**. The integer in these figures shows the number of the intersection of samples belonging to different clusters identified by our method with samples of different clusters identified by xCell. **E**. The distribution of four immune related scores in six immune subtypes identified by xCell. **F**. The differences of overall survival of samples among six immune subtypes identified by xCell.

**Figure S16. The boxplot shows the expression of several key genes in the six immune subtypes across TCGA 33 cancer types**. The p values of variance analysis among six immune subtypes are significant (p < 2e-16).

**Figure S17. Survival analysis of six clusters of 33 cancer samples**.

**Figure S18. The predicted immune cell infiltration by our method based on lncRNA signatures or union sets by combining lncRNA signatures with protein-coding gene signatures from CIBERSORT**.

**Figure S19. The expression of lncRNA signatures in three CD8**^+^ **subtypes**.

**Table S1 .The source of 247 samples**.

**TableS2 . The number and estimation accuracy of lncRNA signature sets**.

**Table S3 . Functional enrichment results (GO)**

**Table S4 . Functional enrichment results (KEGG)**.

## References

[1] M Nuti, I G Zizzari, A Botticelli, A Rughetti, P Marchetti. The ambitious role of anti angiogenesis molecules: Turning a cold tumor into a hot one, Cancer treatment reviews 2018; 70: 41–46.

[2] D M Pardoll. The blockade of immune checkpoints in cancer immunotherapy, Nature reviews. Cancer 2012; 12: 252–64.

[3] T F Gajewski, H Schreiber, Y X Fu. Innate and adaptive immune cells in the tumor microenvironment, Nature immunology 2013; 14: 1014–22.

[4] V Hillerdal, M Essand. Chimeric antigen receptor-engineered T cells for the treatment of metastatic prostate cancer, BioDrugs : clinical immunotherapeutics, biopharmaceuticals and gene therapy 2015; 29: 75–89.

[5] W Hugo, J M Zaretsky, L Sun, C Song, B H Moreno, S Hu-Lieskovan, B Berent-Maoz, J Pang, B Chmielowski, G Cherry, E Seja, S Lomeli, X Kong, M C Kelley, J A Sosman, D B Johnson, A Ribas, R S Lo. Genomic and Transcriptomic Features of Response to Anti-PD-1 Therapy in Metastatic Melanoma, Cell 2016; 165: 35–44.

[6] R Andersen, M Donia, M C Westergaard, M Pedersen, M Hansen, I M Svane. Tumor infiltrating lymphocyte therapy for ovarian cancer and renal cell carcinoma, Human vaccines & immunotherapeutics 2015; 11: 2790–5.

[7] H S Robins, N G Ericson, J Guenthoer, K C O’Briant, M Tewari, C W Drescher, J H Bielas. Digital genomic quantification of tumor-infiltrating lymphocytes, Science translational medicine 2013; 5: 214ra169.

[8] J Galon, A Costes, F Sanchez-Cabo, A Kirilovsky, B Mlecnik, C Lagorce-Pagès, M Tosolini, M Camus, A Berger, P Wind, F Zinzindohoué, P Bruneval, P H Cugnenc, Z Trajanoski, W H Fridman, F Pagès. Type, density, and location of immune cells within human colorectal tumors predict clinical outcome, Science (New York, N.Y.) 2006; 313: 1960–4.

[9] L Ding, J Ren, D Zhang, Y Li, X Huang, Q Hu, H Wang, Y Song, Y Ni, Y Hou. A novel stromal lncRNA signature reprograms fibroblasts to promote the growth of oral squamous cell carcinoma via LncRNA-CAF/interleukin-33, Carcinogenesis 2018; 39: 397–406.

[10] Y Wu, J Li, P Jabbarzadeh Kaboli, J Shen, X Wu, Y Zhao, H Ji, F Du, Y Zhou, Y Wang, H Zhang, J Yin, Q Wen, C H Cho, M Li, Z Xiao. Natural killer cells as a double-edged sword in cancer immunotherapy: A comprehensive review from cytokine therapy to adoptive cell immunotherapy, Pharmacological research 2020; 155: 104691.

[11] J Ji, Y Yin, H Ju, X Xu, W Liu, Q Fu, J Hu, X Zhang, B Sun. Long non-coding RNA Lnc-Tim3 exacerbates CD8 T cell exhaustion via binding to Tim-3 and inducing nuclear translocation of Bat3 in HCC, Cell death & disease 2018; 9: 478.

[12] L Zhao, G Ji, X Le, C Wang, L Xu, M Feng, Y Zhang, H Yang, Y Xuan, Y Yang, L Lei, Q Yang, W B Lau, B Lau, Y Chen, X Deng, S Yao, T Yi, X Zhao, Y Wei, S Zhou. Long Noncoding RNA LINC00092 Acts in Cancer-Associated Fibroblasts to Drive Glycolysis and Progression of Ovarian Cancer, Cancer research 2017; 77: 1369–1382.

[13] S Wang, K Liang, Q Hu, P Li, J Song, Y Yang, J Yao, L S Mangala, C Li, W Yang, P K Park, D H Hawke, J Zhou, Y Zhou, W Xia, M C Hung, J R Marks, G E Gallick, G Lopez-Berestein, E R Flores, A K Sood, S Huang, D Yu, L Yang, C Lin. JAK2-binding long noncoding RNA promotes breast cancer brain metastasis, The Journal of clinical investigation 2017; 127: 4498–4515.

[14] X Pei, X Wang, H Li. LncRNA SNHG1 regulates the differentiation of Treg cells and affects the immune escape of breast cancer via regulating miR-448/IDO, International journal of biological macromolecules 2018; 118: 24–30.

[15] M S Rooney, S A Shukla, C J Wu, G Getz, N Hacohen. Molecular and genetic properties of tumors associated with local immune cytolytic activity, Cell 2015; 160: 48–61.

[16] A M Newman, C L Liu, M R Green, A J Gentles, W Feng, Y Xu, C D Hoang, M Diehn, A A Alizadeh. Robust enumeration of cell subsets from tissue expression profiles, Nature methods 2015; 12: 453–7.

[17] D Aran, Z Hu, A J Butte. xCell: digitally portraying the tissue cellular heterogeneity landscape, Genome biology 2017; 18: 220.

[18] T Li, J Fan, B Wang, N Traugh, Q Chen, J S Liu, B Li, X S Liu. TIMER: a web server for comprehensive analysis of tumor-infiltrating immune cells, Cancer research 2017; 77: e108–e110.

[19] E Becht, N A Giraldo, L Lacroix, B Buttard, N Elarouci, F Petitprez, J Selves, P Laurent-Puig, C Sautès-Fridman, W H Fridman, A de Reyniès. Estimating the population abundance of tissue-infiltrating immune and stromal cell populations using gene expression, Genome biology 2016; 17: 218.

[20] J Racle, K de Jonge, P Baumgaertner, D E Speiser, D Gfeller. Simultaneous enumeration of cancer and immune cell types from bulk tumor gene expression data, eLife 2017; 6.

[21] A Chakravarthy, A Furness, K Joshi, E Ghorani, K Ford, M J Ward, E V King, M Lechner, T Marafioti, S A Quezada, G J Thomas, A Feber, T R Fenton. Pan-cancer deconvolution of tumour composition using DNA methylation, Nature communications 2018; 9: 3220.

[22] T Gong, J D Szustakowski. DeconRNASeq: a statistical framework for deconvolution of heterogeneous tissue samples based on mRNA-Seq data, Bioinformatics (Oxford, England) 2013; 29: 1083–5.

[23] V Ranzani, G Rossetti, I Panzeri, A Arrigoni, R J Bonnal, S Curti, P Gruarin, E Provasi, E Sugliano, M Marconi, R De Francesco, J Geginat, B Bodega, S Abrignani, M Pagani. The long intergenic noncoding RNA landscape of human lymphocytes highlights the regulation of T cell differentiation by linc-MAF-4, Nature immunology 2015; 16: 318–325.

[24] H S Chiu, S Somvanshi, E Patel, T W Chen, V P Singh, B Zorman, S L Patil, Y Pan, S S Chatterjee, A K Sood, P H Gunaratne, P Sumazin. Pan-Cancer Analysis of lncRNA Regulation Supports Their Targeting of Cancer Genes in Each Tumor Context, Cell reports 2018; 23: 297-312.e12.

[25] H Chen, C Li, X Peng, Z Zhou, J N Weinstein, H Liang. A Pan-Cancer Analysis of Enhancer Expression in Nearly 9000 Patient Samples, Cell 2018; 173: 386-399.e12.

[26] C Trapnell, L Pachter, S L Salzberg. TopHat: discovering splice junctions with RNA-Seq, Bioinformatics (Oxford, England) 2009; 25: 1105–11.

[27] C Trapnell, B A Williams, G Pertea, A Mortazavi, G Kwan, M J van Baren, S L Salzberg, B J Wold, L Pachter. Transcript assembly and quantification by RNA-Seq reveals unannotated transcripts and isoform switching during cell differentiation, Nature biotechnology 2010; 28: 511–5.

[28] Z S Bao, H M Chen, M Y Yang, C B Zhang, K Yu, W L Ye, B Q Hu, W Yan, W Zhang, J Akers, V Ramakrishnan, J Li, B Carter, Y W Liu, H M Hu, Z Wang, M Y Li, K Yao, X G Qiu, C S Kang, Y P You, X L Fan, W S Song, R Q Li, X D Su, C C Chen, T Jiang. RNA-seq of 272 gliomas revealed a novel, recurrent PTPRZ1-MET fusion transcript in secondary glioblastomas, Genome research 2014; 24: 1765–73.

[29] X Lin, T Jiang, J Bai, J Li, T Wang, J Xiao, Y Tian, X Jin, T Shao, J Xu, L Chen, L Wang, Y Li. Characterization of Transcriptome Transition Associates Long Noncoding RNAs with Glioma Progression, Molecular therapy. Nucleic acids 2018; 13: 620–632.

[30] S Bhattacharya, P Dunn, C G Thomas, B Smith, H Schaefer, J Chen, Z Hu, K A Zalocusky, R D Shankar, S S Shen-Orr, E Thomson, J Wiser, A J Butte. ImmPort, toward repurposing of open access immunological assay data for translational and clinical research, Scientific data 2018; 5: 180015.

[31] X Zhang, Y Lan, J Xu, F Quan, E Zhao, C Deng, T Luo, L Xu, G Liao, M Yan, Y Ping, F Li, A Shi, J Bai, T Zhao, X Li, Y Xiao. CellMarker: a manually curated resource of cell markers in human and mouse, Nucleic acids research 2019; 47: D721–d728.

[32] M L Telli, D G Stover, S Loi, S Aparicio, L A Carey, S M Domchek, L Newman, G W Sledge, E P Winer. Homologous recombination deficiency and host anti-tumor immunity in triple-negative breast cancer, Breast cancer research and treatment 2018; 171: 21–31.

[33] S Narayanan, T Kawaguchi, L Yan, X Peng, Q Qi, K Takabe. Cytolytic Activity Score to Assess Anticancer Immunity in Colorectal Cancer, Annals of surgical oncology 2018; 25: 2323–2331.

[34] M Lauss, M Donia, K Harbst, R Andersen, S Mitra, F Rosengren, M Salim, J Vallon-Christersson, T Törngren, A Kvist, M Ringnér, I M Svane, G Jönsson. Mutational and putative neoantigen load predict clinical benefit of adoptive T cell therapy in melanoma, Nature communications 2017; 8: 1738.

[35] K Yoshihara, M Shahmoradgoli, E Martínez, R Vegesna, H Kim, W Torres-Garcia, V Treviño, H Shen, P W Laird, D A Levine, S L Carter, G Getz, K Stemke-Hale, G B Mills, R G Verhaak. Inferring tumour purity and stromal and immune cell admixture from expression data, Nature communications 2013; 4: 2612.

[36] J M Engreitz, K Sirokman, P McDonel, A A Shishkin, C Surka, P Russell, S R Grossman, A Y Chow, M Guttman, E S Lander. RNA-RNA interactions enable specific targeting of noncoding RNAs to nascent Pre-mRNAs and chromatin sites, Cell 2014; 159: 188–199.

[37] D K Shah, J C Zúñiga-Pflücker. An overview of the intrathymic intricacies of T cell development, Journal of immunology (Baltimore, Md. : 1950) 2014; 192: 4017–23.

[38] J W Pollard. Trophic macrophages in development and disease, Nature reviews. Immunology 2009; 9: 259–70.

[39] Y X Zhou, W Zhao, L W Mao, Y L Wang, L Q Xia, M Cao, J Shen, J Chen. Long non-coding RNA NIFK-AS1 inhibits M2 polarization of macrophages in endometrial cancer through targeting miR-146a, The international journal of biochemistry & cell biology 2018; 104: 25–33.

[40] B J Schmiedel, D Singh, A Madrigal, A G Valdovino-Gonzalez, B M White, J Zapardiel-Gonzalo, B Ha, G Altay, J A Greenbaum, G McVicker, G Seumois, A Rao, M Kronenberg, B Peters, P Vijayanand. Impact of Genetic Polymorphisms on Human Immune Cell Gene Expression, Cell 2018; 175: 1701-1715.e16.

[41] W K Mowel, J J Kotzin, S J McCright, V D Neal, J Henao-Mejia. Control of Immune Cell Homeostasis and Function by lncRNAs, Trends in immunology 2018; 39: 55–69.

[42] X Hu, S Goswami, J Qiu, Q Chen, S Laverdure, B T Sherman, T Imamichi. Profiles of Long Non-Coding RNAs and mRNA Expression in Human Macrophages Regulated by Interleukin-27, International journal of molecular sciences 2019; 20.

[43] L L Liu, S G Zhu, X Y Jiang, J Ren, Y Lin, N N Zhang, M L Tong, H L Zhang, W H Zheng, H J Fu, H J Luo, L R Lin, J H Yan, T C Yang. LncRNA Expression in CD4+ T Cells in Neurosyphilis Patients, Frontiers in cellular and infection microbiology 2017; 7: 461.

[44] D Huang, J Chen, L Yang, Q Ouyang, J Li, L Lao, J Zhao, J Liu, Y Lu, Y Xing, F Chen, F Su, H Yao, Q Liu, S Su, E Song. NKILA lncRNA promotes tumor immune evasion by sensitizing T cells to activation-induced cell death, Nature immunology 2018; 19: 1112–1125.

[45] D Zemmour, A Pratama, S M Loughhead, D Mathis, C Benoist. Flicr, a long noncoding RNA, modulates Foxp3 expression and autoimmunity, Proceedings of the National Academy of Sciences of the United States of America 2017; 114: E3472–e3480.

[46] A Mantovani, F Marchesi, A Malesci, L Laghi, P Allavena. Tumour-associated macrophages as treatment targets in oncology, Nature reviews. Clinical oncology 2017; 14: 399–416.

[47] A Gieryng, D Pszczolkowska, K A Walentynowicz, W D Rajan, B Kaminska. Immune microenvironment of gliomas, Laboratory investigation; a journal of technical methods and pathology 2017; 97: 498–518.

[48] P Domingues, M González-Tablas, Á Otero, D Pascual, D Miranda, L Ruiz, P Sousa, J Ciudad, J M Gonçalves, M C Lopes, A Orfao, M D Tabernero. Tumor infiltrating immune cells in gliomas and meningiomas, Brain, behavior, and immunity 2016; 53: 1–15.

[49] G L Lin, S Nagaraja, M G Filbin, M L Suvà, H Vogel, M Monje. Non-inflammatory tumor microenvironment of diffuse intrinsic pontine glioma, Acta neuropathologica communications 2018; 6: 51.

[50] N A P Lieberman, K DeGolier, H M Kovar, A Davis, V Hoglund, J Stevens, C Winter, G Deutsch, S N Furlan, N A Vitanza, S E S Leary, C A Crane. Characterization of the immune microenvironment of diffuse intrinsic pontine glioma: implications for development of immunotherapy, Neuro-oncology 2019; 21: 83–94.

[51] C Zhang, W Cheng, X Ren, Z Wang, X Liu, G Li, S Han, T Jiang, A Wu. Tumor Purity as an Underlying Key Factor in Glioma, Clinical cancer research : an official journal of the American Association for Cancer Research 2017; 23: 6279–6291.

[52] F S Varn, Y Wang, D W Mullins, S Fiering, C Cheng. Systematic Pan-Cancer Analysis Reveals Immune Cell Interactions in the Tumor Microenvironment, Cancer research 2017; 77: 1271–1282.

[53] X Liu, S Wu, Y Yang, M Zhao, G Zhu, Z Hou. The prognostic landscape of tumor-infiltrating immune cell and immunomodulators in lung cancer, Biomedicine & pharmacotherapy = Biomedecine & pharmacotherapie 2017; 95: 55–61.

[54] J Peng, H Xu, Y Chen, W Wang, L Zhu, Y Shao, J Wang. Screening for therapeutic targets of tumor angiogenesis signatures in 31 cancer types and potential insights, Biochemical and biophysical research communications 2019; 508: 465–471.

[55] H Rahimi Koshkaki, S Minasi, A Ugolini, G Trevisi, C Napoletano, I G Zizzari, M Gessi, F Giangaspero, A Mangiola, M Nuti, F R Buttarelli, A Rughetti. Immunohistochemical Characterization of Immune Infiltrate in Tumor Microenvironment of Glioblastoma, Journal of personalized medicine 2020; 10.

[56] R Zhong, D Chen, S Cao, J Li, B Han, H Zhong. Immune cell infiltration features and related marker genes in lung cancer based on single-cell RNA-seq, Clinical & translational oncology : official publication of the Federation of Spanish Oncology Societies and of the National Cancer Institute of Mexico 2021; 23: 405–417.

[57] J Cursons, F Souza-Fonseca-Guimaraes, M Foroutan, A Anderson, F Hollande, S Hediyeh-Zadeh, A Behren, N D Huntington, M J Davis. A Gene Signature Predicting Natural Killer Cell Infiltration and Improved Survival in Melanoma Patients, Cancer immunology research 2019; 7: 1162–1174.

[58] J Lan, L Sun, F Xu, L Liu, F Hu, D Song, Z Hou, W Wu, X Luo, J Wang, X Yuan, J Hu, G Wang. M2 Macrophage-Derived Exosomes Promote Cell Migration and Invasion in Colon Cancer, Cancer research 2019; 79: 146–158.

[59] X Wang, G Luo, K Zhang, J Cao, C Huang, T Jiang, B Liu, L Su, Z Qiu. Hypoxic Tumor-Derived Exosomal miR-301a Mediates M2 Macrophage Polarization via PTEN/PI3Kγ to Promote Pancreatic Cancer Metastasis, Cancer research 2018; 78: 4586–4598.

[60] C Rodríguez-Cerdeira, M Carnero Gregorio, A López-Barcenas, E Sánchez-Blanco, B Sánchez-Blanco, G Fabbrocini, B Bardhi, A Sinani, R A Guzman. Advances in Immunotherapy for Melanoma: A Comprehensive Review, Mediators of inflammation 2017; 2017: 3264217.

[61] P Danaher, S Warren, L Dennis, L D’Amico, A White, M L Disis, M A Geller, K Odunsi, J Beechem, S P Fling. Gene expression markers of Tumor Infiltrating Leukocytes, Journal for immunotherapy of cancer 2017; 5: 18.

[62] L E Kandalaft, D J Powell, Jr., N Singh, G Coukos. Immunotherapy for ovarian cancer: what’s next?, Journal of clinical oncology : official journal of the American Society of Clinical Oncology 2011; 29: 925–33.

[63] H Schuster, J K Peper, H C Bösmüller, K Röhle, L Backert, T Bilich, B Ney, M W Löffler, D J Kowalewski, N Trautwein, A Rabsteyn, T Engler, S Braun, S P Haen, J S Walz, B Schmid-Horch, S Y Brucker, D Wallwiener, O Kohlbacher, F Fend, H G Rammensee, S Stevanović, A Staebler,P Wagner. The immunopeptidomic landscape of ovarian carcinomas, Proceedings of the National Academy of Sciences of the United States of America 2017; 114: E9942–e9951.

[64] R D Schreiber, L J Old,M J Smyth. Cancer immunoediting: integrating immunity’s roles in cancer suppression and promotion, Science (New York, N.Y.) 2011; 331: 1565–70.

[65] L J Kinlen. Immunosuppression and cancer, IARC scientific publications 1992; 237–53.

[66] G P Dunn, A T Bruce, H Ikeda, L J Old,R D Schreiber. Cancer immunoediting: from immunosurveillance to tumor escape, Nature immunology 2002; 3: 991–8.

[67] T X Huang, L Fu. The immune landscape of esophageal cancer, Cancer communications (London, England) 2019; 39: 79.

[68] M Swart, I Verbrugge, J B Beltman. Combination Approaches with Immune-Checkpoint Blockade in Cancer Therapy, Frontiers in oncology 2016; 6: 233.

[69] P Jiang, S Gu, D Pan, J Fu, A Sahu, X Hu, Z Li, N Traugh, X Bu, B Li, J Liu, G J Freeman, M A Brown, K W Wucherpfennig, X S Liu. Signatures of T cell dysfunction and exclusion predict cancer immunotherapy response, Nature medicine 2018; 24: 1550–1558.

[70] M B Schaaf, A D Garg, P Agostinis. Defining the role of the tumor vasculature in antitumor immunity and immunotherapy, Cell death & disease 2018; 9: 115.

[71] Y R Miao, Q Zhang, Q Lei, M Luo, G Y Xie, H Wang, A Y Guo. ImmuCellAI: A Unique Method for Comprehensive T-Cell Subsets Abundance Prediction and its Application in Cancer Immunotherapy, Advanced science (Weinheim, Baden-Wurttemberg, Germany) 2020; 7: 1902880.

[72] D S Vinay, E P Ryan, G Pawelec, W H Talib, J Stagg, E Elkord, T Lichtor, W K Decker, R L Whelan, H Kumara, E Signori, K Honoki, A G Georgakilas, A Amin, W G Helferich, C S Boosani, G Guha, M R Ciriolo, S Chen, S I Mohammed, A S Azmi, W N Keith, A Bilsland, D Bhakta, D Halicka, H Fujii, K Aquilano, S S Ashraf, S Nowsheen, X Yang, B K Choi,B S Kwon. Immune evasion in cancer: Mechanistic basis and therapeutic strategies, Seminars in cancer biology 2015; 35 Suppl: S185–s198.

